# TMSNP: a web server to predict pathogenesis of missense mutations in transmembrane region of membrane proteins

**DOI:** 10.1101/2020.01.21.913764

**Authors:** Adrián Garcia-Recio, José Carlos Gómez-Tamayo, Iker Reina, Mercedes Campillo, Arnau Cordomí, Mireia Olivella

## Abstract

The massive amount of data generated from genome sequencing have given rise to several mutation predictor tools although no mutation database or predictor tool have been developed specifically for the transmembrane region of membrane proteins.

We present TMSNP, a database that currently contains information from 2624 pathogenic and 195964 non-pathogenic reported mutations located on the TM region of membrane proteins. The computed conservation parameters and annotations on these mutations were used to train a machine-learning model that classifies TM mutations as pathogenic or non-pathogenic. The presented tool improves considerably the prediction power of commonly used mutation predictors and additionally represents the first mutation prediction tool specific for TM mutations.

TMSNP is available at http://lmc.uab.es/tmsnp/

## Introduction

Membrane proteins represent 25% of all human proteins (Dobson, et al., 2015; Gromiha and Ou, 2014) and perform essential roles in cellular functions. Approximately 50-60% of TM proteins are drug targets for various diseases (Almeida, et al., 2017; Overington, et al., 2006) and 90% of membrane proteins present disease-associated missense mutations that may affect protein folding, stability and function (Kulandaisamy, et al., 2019). Whole genome and exome sequencing have revealed that missense mutations that are mendelian and rare disease-causing are more frequent than previously thought and collectively affect millions of patients worldwide (Chong, et al., 2015). Thus, there is an urgent need to understand the relation between genotype and phenotype in order to identify disease causing genetic variants within candidate variants. Variant prioritization tools such as SIFT (Sim, et al., 2012) or Polyphen-2 (Adzhubei, et al., 2010) are widely used to predict the effect of mutations based on evolutionary conservation and expected impact on structure and function. Although the transmembrane region of membrane proteins differs from globular proteins in terms of sequence-structure conservation (Olivella, et al., 2013), amino acid distribution and inter-residue interactions (Mayol, et al., 2018), no mutation prediction tool have been specifically developed for the transmembrane region of membrane proteins. Here we present TMSNP (accessible at http://lmc.uab.es/tmsnp/), a database and a mutation predictor server trained and adapted to the specific features of transmembrane proteins using the information of the position and amino acid change encoded in Pfam alignments to predict pathogenicity of TM missense mutations.

## Methods

We took i) all human membrane proteins tagged as reviewed, ii) the ranges of residues that form TM segments, and iii) position annotations from the Uniprot (McGarvey, et al., 2019; UniProt, 2019). Pathogenic missense mutations located in these human TM segments were taken from ClinVar (Landrum, et al., 2014) and SwissVar (Mottaz, et al., 2010), and only those annotated as disease-causing/pathogenic for a mendelian disorder were kept. We also retrieved non-pathogenic missense mutations and its population allele frequency from GnomAD (Karczewski, et al., 2019). The obtained pathogenic and non-pathogenic missense mutations were used to classify human TM proteins as “pathogenic proteins”, when at least one disease-causing pathogenic mutation has been reported for this protein and as “non-pathogenic proteins”, elsewhere. Multiple sequence alignments of all human TM domains were taken from Pfam database (El-Gebali, et al., 2019). For each missense mutation we computed the following parameters: (i) amino acid type and frequencies of the wild type (wt) and mutated amino acids, (ii) score associated to wt/mutant amino acid change in the PHAT 75/73 substitution matrix, specific for membrane proteins (Ng, et al., 2000) and (iii) the entropy of the information content (Pei and Grishin, 2001).

We discarded missense mutations in proteins for which no pathogenic disease-causing mutations have been reported, those likely involved in complex diseases or recessive inheritance (Eilbeck, et al., 2017). This permitted to select missense mutations whose pathogenicity can be linked to protein function and/or structure alteration. We performed homology reduction by discarding homologous mutations in the same position in the Pfam alignment. The filtered dataset contained 2704 pathogenic and 19292 non-pathogenic TM mutations and was subsequently subsampled to obtain a balanced dataset. This was done by selecting non-pathogenic mutations with the highest population allele frequency from GnomAD (Karczewski, et al., 2019), to ensure that these were neutral mutations. The resulting balanced dataset matrix had 5408 missense mutations (50% pathogenic and 50% non-pathogenic) and 569 variables (***569V dataset***; see http://lmc.uab.es/tmsnp/569Vdataset). A subset from the 569V dataset that only uses the four most contributive variables associated to conservation (see Supp Table 1): initial frequency, final frequency, matrix score and entropy was also constructed (***4V dataset***). Thus, the Uniprot accession code and the Pfams code variables were not used in this dataset. For each Uniprot code in the 569V and 4V dataset, mutations were split in a training set (80%) that was used to build machine-learning models, while the remaining 20% of the samples were used in external-validation. Although homologous mutations (i.e. same amino acid change in the same Pfam alignment position) were previously excluded in the 569V and 4V dataset, we wanted to exclude any possible bias due to the presence of homologous proteins in the validation set. Using the 569V dataset, mutations with the same Pfam code were used exclusively in either the training (80%) or the validation set (20%) (***569V Exclusive Pfams dataset***). Consequently, mutations with the same Uniprot code were also used in either the training set or validation set.

**Table 1.** Sensitivity, specificity, Matthews correlation coefficient (MCC) and coverage of TMSNP model (569V dataset) at various levels of significance in external validation. Comparison to SIFT and Poyphen-2 is also presented. MCC is a quality metric which rewards models with balanced sensitivity and specificity. Coverage stands for the percentage of samples inside the applicability domain.

| Method | Sensitivity | Specificity | MCC | Coverage |
| --- | --- | --- | --- | --- |
| **TMSNP**<br>(0.05 significance) | 0.92 | 0.87 | 0.80 | 0.43 |
| **TMSNP**<br>(0.1 significance) | 0.89 | 0.82 | 0.72 | 0.63 |
| **TMSNP**<br>(0.2 significance) | 0.83 | 0.76 | 0.59 | 0.89 |
| **Polyhen-2** | 0.93 | 0.35 | 0.64 | 1 |
| **SIFT** | 0.88 | 0.52 | 0.70 | 1 |

Machine-learning models were built in Python 3 using Flame modeling framework (https://github.com/phi-grib/flame), which is based on Scikit-learn library (http://scikit-learn.sourceforge.net). We used conformal prediction framework as applicability domain technique (Norinder, et al., 2014). Various predictive models using different settings on the algorithm, applicability domain, and dataset were built and were internally validated using K-fold (K=5) cross-validation.

TMSNP web application tool was constructed on a Python 3.6.6 backend with the Flask 1.0.2 framework (http://flask.pocoo.org). TMSMP and the corresponding datasets used for training and testing the predictor are compiled automatically using Python scripts that access data from Databases in a MySQL database, thus facilitating regular updates.

## Results

TMSNP currently contains a database of 2704 pathogenic and 192566 non-pathogenic mutations located in the TM segments of human membrane proteins (see Figure 1). Pathogenic and non-pathogenic mutations in disease associated membrane proteins were used to develop an algorithm in machine learning models using Random Forest (Supp. Table 2 and Supp. Table 3). The three models (569V, 4V, 569V exclusive PFAMs) show excellent performance, although the 569V dataset increases both the quality statistics and the confidence in predictions (reflected in the higher coverage). The small loss in accuracy in the two other models is attributed to the contribution to propensity to pathogenicity for each protein, which is associated to the Uniprot code. We chose to implement Random Forest algorithm on 569V dataset because of better results in external validation. Because better significance comes at a cost of a lower coverage, for a given SNP, TMSNP returns the unambiguous class prediction at the higher confidence possible. The predictions with a confidence below 0.25 are considered outside the domain of applicability. Table 1 shows the comparison between TMSNP models (569V dataset) generated at three levels of significance and SIFT and Polyphen-2 mutation prediction tools. Lower significance at the conformal predictor increases the performance at a cost of lower coverage. When compared to SIFT and Polyphen-2, the prediction power of TMSNP for TM missense mutations has similar sensitivity but remarkably higher specificity, resulting in a significant predictive power improvement reflected in the Matthews correlation coefficient value (Russell 2012).

**Figure 1.**
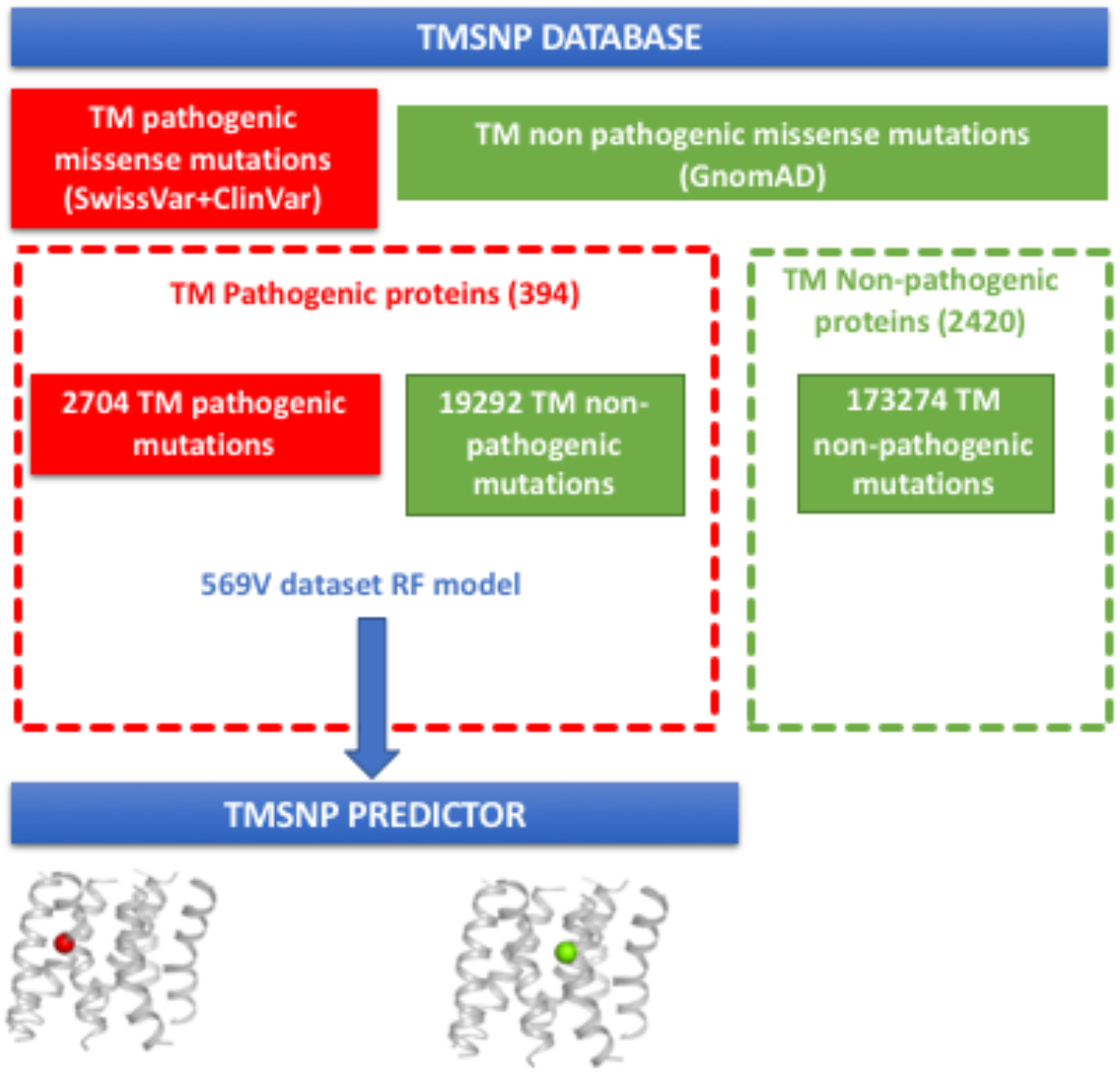
TMSNP recopilates pathogenic and non-pathogenic missense mutations in TM segments of membrane proteins from SwissVar, Uniprot and GnomAD. Proteins are split into i) proteins without any reported disease-causing mutation and ii) proteins with reported disease-causing mutation. 569 features in pathogenic proteins are trained by machine learning methods to develop TMSNP prediction model that predicts pathogenicity of TM mutations.

TMSNP is a free regularly updated web server that presents two main functionalities: (i) a database of reported pathogenic and non-pathogenic mutations in TM segments of membrane proteins (ii) a mutation prediction tool able to predict pathogenicity and its confidence interval for previously non-reported TM missense mutations. The prediction algorithm developed specifically for membrane proteins allows to considerably improve the prediction power compared to current mutation predictor servers.

