## Supplementary Material for "TMSNP: a web server to predict pathogenesis of missense mutations in transmembrane region of membrane proteins"

**Supplementary Table 1**. Contribution of the 569 features to the Random Forest algorithm.

| *Feature* | *Contribution* |
|---|---|
| *Final frequency* | *10.4%* |
| *Entropy* | *8.3%* |
| *Initial frequency* | *8.3%* |
| *Matrix Score* | *5.4%* |
| *Uniprot codes global contribution (358 features)* | *31.9%* |
| *Pfam codes global contribution (167 features)* | *18.6%* |
| *Reference amino acid global contribution (20 features)* | *7.5%* |
| *Mutated amino acid global contribution (20 features)* | *8.9%* |

**Supplementary Table 2.** Model statistics in cross-validation. Quality metrics in cross-validation (5-fold) for the machine-learning models created using three datasets 569V, 4V, 569V with exclusive Pfams with different conformal significance. Replica refers to the unique model using a determined random seed for sampling into train and test sets . TP accounts for true positives, TN for true negatives, FP for false positives and FN for false negatives. MCC stands for Matthews Correlation coefficient, which is a measure that combines Sensitivity and Specificity. Coverage stands for the percentage of samples inside the applicability domain.

| Dataset | Significance | Replica | TP | TN | FP | FN | Sensitivity | Specificity | MCC | Coverage | Accuracy |
|---|---|---|---|---|---|---|---|---|---|---|---|
| **569V dataset** | **0.05** | 1 | 810 | 691 | 89 | 82 | 0.91 | 0.89 | 0.79 | 0.39 | 0.90 |
| | | 2 | 874 | 674 | 94 | 89 | 0.91 | 0.88 | 0.79 | 0.40 | 0.89 |
| | | 3 | 851 | 717 | 97 | 89 | 0.91 | 0.88 | 0.79 | 0.41 | 0.89 |
| | | 4 | 911 | 721 | 113 | 99 | 0.90 | 0.87 | 0.77 | 0.43 | 0.89 |
| | | 5 | 855 | 683 | 116 | 80 | 0.91 | 0.86 | 0.77 | 0.40 | 0.89 |

| | | | | | | | | | | | |
|---|---|---|---|---|---|---|---|---|---|---|---|
| | | **mean** | | | | | **0.91** | **0.87** | **0.78** | **0.40** | **0.89** |
| | | SD | | | | | 0.01 | 0.01 | 0.01 | 0.01 | 0.01 |
| | **0.1** | 1 | 810 | 1001 | 215 | 168 | 0.88 | 0.82 | 0.70 | 0.60 | 0.85 |
| | | 2 | 1230 | 988 | 212 | 172 | 0.88 | 0.82 | 0.70 | 0.60 | 0.85 |
| | | 3 | 1189 | 1034 | 200 | 176 | 0.87 | 0.84 | 0.71 | 0.60 | 0.86 |
| | | 4 | 1246 | 1027 | 210 | 192 | 0.87 | 0.83 | 0.70 | 0.62 | 0.85 |
| | | 5 | 1208 | 977 | 216 | 174 | 0.87 | 0.82 | 0.70 | 0.60 | 0.85 |
| | | **mean** | | | | | **0.87** | **0.83** | **0.70** | **0.60** | **0.85** |
| | | SD | | | | | 0.01 | 0.01 | 0.01 | 0.01 | 0.00 |
| | **0.2** | 1 | 1584 | 1384 | 437 | 392 | 0.80 | 0.76 | 0.56 | 0.88 | 0.78 |
| | | 2 | 1611 | 1381 | 423 | 375 | 0.81 | 0.77 | 0.58 | 0.88 | 0.79 |
| | | 3 | 1541 | 1460 | 410 | 397 | 0.80 | 0.78 | 0.58 | 0.88 | 0.79 |
| | | 4 | 1605 | 1391 | 450 | 391 | 0.80 | 0.76 | 0.56 | 0.89 | 0.78 |
| | | 5 | 1603 | 1347 | 456 | 376 | 0.81 | 0.75 | 0.56 | 0.87 | 0.78 |
| | | **mean** | | | | | **0.80** | **0.76** | **0.57** | **0.88** | **0.78** |
| | | SD | | | | | 0.007 | 0.013 | 0.009 | 0.005 | 0.004 |
| **4V dataset** | **0.05** | 1 | 258 | 112 | 82 | 28 | 0.90 | 0.58 | 0.52 | 0.11 | 0.77 |
| | | 2 | 314 | 130 | 90 | 43 | 0.88 | 0.59 | 0.50 | 0.13 | 0.77 |
| | | 3 | 245 | 146 | 81 | 60 | 0.80 | 0.64 | 0.45 | 0.12 | 0.74 |
| | | 4 | 304 | 124 | 99 | 77 | 0.80 | 0.56 | 0.36 | 0.14 | 0.71 |
| | | 5 | 290 | 135 | 85 | 46 | 0.86 | 0.61 | 0.50 | 0.13 | 0.76 |
| | | **mean** | | | | | **0.85** | **0.60** | **0.47** | **0.13** | **0.75** |
| | | SD | | | | | 0.047 | 0.034 | 0.062 | 0.011 | 0.027 |
| | **0.1** | 1 | 503 | 352 | 201 | 118 | 0.81 | 0.64 | 0.46 | 0.27 | 0.73 |

| | | | | | | | | | | | |
|---|---|---|---|---|---|---|---|---|---|---|---|
| | | 2 | 556 | 378 | 177 | 125 | 0.82 | 0.68 | 0.50 | 0.29 | 0.76 |
| | | 3 | 488 | 386 | 166 | 169 | 0.74 | 0.70 | 0.44 | 0.28 | 0.72 |
| | | 4 | 575 | 362 | 202 | 171 | 0.77 | 0.64 | 0.42 | 0.30 | 0.72 |
| | | 5 | 540 | 368 | 191 | 138 | 0.80 | 0.66 | 0.46 | 0.29 | 0.73 |
| | | **mean** | | | | | **0.79** | **0.66** | **0.46** | **0.29** | **0.73** |
| | | SD | | | | | 0.030 | 0.027 | 0.032 | 0.012 | 0.015 |
| | **0.2** | 1 | 887 | 776 | 391 | 351 | 0.72 | 0.67 | 0.38 | 0.56 | 0.69 |
| | | 2 | 947 | 767 | 389 | 385 | 0.71 | 0.66 | 0.38 | 0.58 | 0.69 |
| | | 3 | 839 | 779 | 358 | 401 | 0.68 | 0.69 | 0.36 | 0.55 | 0.68 |
| | | 4 | 925 | 777 | 401 | 374 | 0.71 | 0.66 | 0.37 | 0.57 | 0.69 |
| | | 5 | 916 | 752 | 403 | 358 | 0.72 | 0.65 | 0.37 | 0.56 | 0.69 |
| | | **mean** | | | | | **0.71** | **0.67** | **0.37** | **0.56** | **0.69** |
| | | SD | | | | | 0.017 | 0.013 | 0.007 | 0.011 | 0.004 |
| **569V Exclusive Pfam Dataset** | **0.05** | 1 | 1049 | 777 | 99 | 95 | 0.92 | 0.89 | 0.80 | 0.43 | 0.90 |
| | | 2 | 1076 | 851 | 113 | 90 | 0.92 | 0.88 | 0.81 | 0.44 | 0.91 |
| | | 3 | 1075 | 862 | 102 | 96 | 0.92 | 0.89 | 0.81 | 0.45 | 0.91 |
| | | 4 | 1265 | 638 | 115 | 78 | 0.94 | 0.85 | 0.80 | 0.45 | 0.91 |
| | | 5 | 1159 | 811 | 120 | 91 | 0.93 | 0.87 | 0.80 | 0.46 | 0.90 |
| | | **mean** | | | | | **0.93** | **0.88** | **0.81** | **0.45** | **0.91** |
| | | SD | | | | | 0.010 | 0.018 | 0.005 | 0.012 | 0.002 |
| | **0.1** | 1 | 1406 | 1139 | 228 | 194 | 0.88 | 0.83 | 0.71 | 0.63 | 0.86 |
| | | 2 | 1477 | 1214 | 236 | 196 | 0.88 | 0.84 | 0.72 | 0.65 | 0.86 |
| | | 3 | 1469 | 1224 | 224 | 207 | 0.88 | 0.85 | 0.72 | 0.66 | 0.86 |
| | | 4 | 1663 | 980 | 238 | 175 | 0.91 | 0.81 | 0.72 | 0.65 | 0.87 |

| | | | | | | | | | | | |
|---|---|---|---|---|---|---|---|---|---|---|---|
| | | 5 | 1555 | 1181 | 253 | 196 | 0.89 | 0.82 | 0.72 | 0.68 | 0.86 |
| | | **mean** | | | | | **0.89** | **0.83** | **0.72** | **0.65** | **0.86** |
| | | SD | | | | | 0.011 | 0.016 | 0.004 | 0.017 | 0.003 |
| | **0.2** | 1 | 1806 | 1573 | 467 | 420 | 0.81 | 0.77 | 0.58 | 0.91 | 0.79 |
| | | 2 | 1850 | 1663 | 477 | 426 | 0.81 | 0.78 | 0.59 | 0.92 | 0.80 |
| | | 3 | 1837 | 1654 | 480 | 417 | 0.82 | 0.78 | 0.59 | 0.93 | 0.80 |
| | | 4 | 2063 | 1405 | 503 | 369 | 0.85 | 0.74 | 0.59 | 0.92 | 0.80 |
| | | 5 | 1929 | 1589 | 475 | 417 | 0.82 | 0.77 | 0.59 | 0.93 | 0.80 |
| | | **mean** | | | | | **0.82** | **0.77** | **0.59** | **0.92** | **0.80** |
| | | SD | | | | | 0.015 | 0.017 | 0.004 | 0.011 | 0.003 |

**Supplementary Table 3**. Model statistics at external-validation. Table shows performance metrics in external-validation (20% of the original dataset) for the machine-learning models created using three datasets (569V, 4V, 569V with exclusive Pfams) with different conformal significance. Replica refers to the unique model using a determined random seed for sampling into train and test sets. TP accounts for true positives, TN for true negatives, FP for false positives and FN for false negatives. MCC accounts for Matthews correlation coefficient (MCC), which is a measure that combines sensitivity and specificity. Coverage stands for the percentage of samples inside the applicability domain.

| Dataset | Significance | Replica | TP | TN | FP | FN | Sensitivity | Specificity | MCC | Coverage | Accuracy |
|---|---|---|---|---|---|---|---|---|---|---|---|
| **569V dataset** | **0.05** | 1 | 212 | 212 | 23 | 23 | 0.90 | 0.90 | 0.80 | 0.44 | 0.90 |
| | | 2 | 195 | 208 | 37 | 14 | 0.93 | 0.85 | 0.78 | 0.42 | 0.89 |
| | | 3 | 217 | 194 | 29 | 21 | 0.91 | 0.87 | 0.78 | 0.43 | 0.89 |
| | | 4 | 209 | 205 | 37 | 13 | 0.94 | 0.85 | 0.79 | 0.43 | 0.89 |
| | | 5 | 223 | 192 | 25 | 16 | 0.93 | 0.89 | 0.82 | 0.42 | 0.91 |
| | | **mean** | | | | | **0.92** | **0.87** | **0.80** | **0.43** | **0.90** |
| | | SD | | | | | 0.017 | 0.024 | 0.017 | 0.006 | 0.009 |
| | **0.1** | 6 | 290 | 294 | 54 | 36 | 0.89 | 0.85 | 0.73 | 0.62 | 0.87 |
| | | 7 | 279 | 308 | 67 | 35 | 0.89 | 0.82 | 0.71 | 0.64 | 0.85 |
| | | 8 | 304 | 286 | 72 | 48 | 0.86 | 0.80 | 0.66 | 0.66 | 0.83 |
| | | 9 | 284 | 290 | 66 | 32 | 0.90 | 0.82 | 0.71 | 0.62 | 0.85 |
| | | 10 | 303 | 288 | 58 | 25 | 0.92 | 0.83 | 0.76 | 0.62 | 0.88 |
| | | **mean** | | | | | **0.89** | **0.82** | **0.72** | **0.63** | **0.86** |
| | | SD | | | | | 0.022 | 0.017 | 0.035 | 0.015 | 0.017 |
| | **0.2** | 11 | 366 | 394 | 119 | 81 | 0.82 | 0.77 | 0.59 | 0.89 | 0.79 |
| | | 12 | 350 | 418 | 128 | 75 | 0.82 | 0.77 | 0.59 | 0.90 | 0.79 |
| | | 13 | 389 | 383 | 131 | 80 | 0.83 | 0.75 | 0.58 | 0.91 | 0.79 |
| | | 14 | 357 | 397 | 136 | 80 | 0.82 | 0.75 | 0.56 | 0.90 | 0.78 |
| | | 15 | 374 | 394 | 110 | 61 | 0.86 | 0.78 | 0.64 | 0.87 | 0.82 |
| | | **mean** | | | | | **0.83** | **0.76** | **0.59** | **0.89** | **0.79** |

| | | | | | | | | | | | |
|---|---|---|---|---|---|---|---|---|---|---|---|
| | | SD | | | | | 0.017 | 0.016 | 0.031 | 0.015 | 0.015 |
| **4V dataset** | **0.05** | 1 | 52 | 48 | 20 | 9 | 0.85 | 0.71 | 0.56 | 0.12 | 0.78 |
| | | 2 | 53 | 40 | 34 | 9 | 0.86 | 0.54 | 0.41 | 0.13 | 0.68 |
| | | 3 | 56 | 34 | 23 | 10 | 0.85 | 0.60 | 0.46 | 0.11 | 0.73 |
| | | 4 | 55 | 40 | 31 | 7 | 0.89 | 0.56 | 0.47 | 0.12 | 0.71 |
| | | 5 | 65 | 36 | 31 | 11 | 0.86 | 0.54 | 0.42 | 0.13 | 0.71 |
| | | **mean** | | | | | **0.86** | **0.59** | **0.46** | **0.12** | **0.72** |
| | | SD | | | | | 0.016 | 0.070 | 0.060 | 0.007 | 0.034 |
| | **0.1** | 6 | 116 | 128 | 43 | 36 | 0.76 | 0.75 | 0.51 | 0.30 | 0.76 |
| | | 7 | 108 | 111 | 64 | 34 | 0.76 | 0.63 | 0.39 | 0.29 | 0.69 |
| | | 8 | 123 | 94 | 58 | 41 | 0.75 | 0.62 | 0.37 | 0.29 | 0.69 |
| | | 9 | 112 | 101 | 62 | 30 | 0.79 | 0.62 | 0.41 | 0.28 | 0.70 |
| | | 10 | 119 | 94 | 63 | 29 | 0.80 | 0.60 | 0.41 | 0.28 | 0.70 |
| | | **mean** | | | | | 0.77 | 0.64 | 0.42 | 0.29 | 0.71 |
| | | SD | | | | | 0.022 | 0.060 | 0.053 | 0.007 | 0.028 |
| | **0.2** | 11 | 199 | 234 | 117 | 86 | 0.70 | 0.67 | 0.36 | 0.59 | 0.68 |
| | | 12 | 183 | 218 | 131 | 90 | 0.67 | 0.63 | 0.29 | 0.58 | 0.65 |
| | | 13 | 219 | 197 | 120 | 87 | 0.72 | 0.62 | 0.34 | 0.58 | 0.67 |
| | | 14 | 196 | 198 | 118 | 91 | 0.68 | 0.63 | 0.31 | 0.56 | 0.65 |
| | | 15 | 201 | 202 | 127 | 65 | 0.76 | 0.61 | 0.37 | 0.55 | 0.68 |
| | | **mean** | | | | | **0.71** | **0.63** | **0.34** | **0.57** | **0.67** |
| | | SD | | | | | 0.033 | 0.021 | 0.033 | 0.015 | 0.015 |
| **569V Exclusive Pfam Dataset** | **0.05** | 1 | 174 | 245 | 35 | 38 | 0.82 | 0.88 | 0.70 | 0.42 | 0.85 |
| | | 2 | 164 | 248 | 34 | 46 | 0.78 | 0.88 | 0.67 | 0.45 | 0.84 |

| | | | | | | | | | | | |
|---|---|---|---|---|---|---|---|---|---|---|---|
| | | 3 | 147 | 286 | 31 | 95 | 0.61 | 0.90 | 0.54 | 0.48 | 0.78 |
| | | 4 | 151 | 290 | 69 | 24 | 0.86 | 0.81 | 0.64 | 0.46 | 0.83 |
| | | 5 | 150 | 316 | 39 | 60 | 0.71 | 0.89 | 0.62 | 0.49 | 0.83 |
| | | **mean** | | | | | **0.76** | **0.87** | **0.63** | **0.46** | **0.82** |
| | | SD | | | | | 0.100 | 0.037 | 0.059 | 0.027 | 0.029 |
| | **0.1** | 6 | 265 | 326 | 73 | 74 | 0.78 | 0.82 | 0.60 | 0.63 | 0.80 |
| | | 7 | 229 | 337 | 58 | 106 | 0.68 | 0.85 | 0.55 | 0.67 | 0.78 |
| | | 8 | 222 | 369 | 69 | 147 | 0.60 | 0.84 | 0.46 | 0.70 | 0.73 |
| | | 9 | 206 | 396 | 110 | 46 | 0.82 | 0.78 | 0.57 | 0.65 | 0.79 |
| | | 10 | 227 | 395 | 76 | 107 | 0.68 | 0.84 | 0.53 | 0.69 | 0.77 |
| | | **mean** | | | | | **0.71** | **0.83** | **0.54** | **0.67** | **0.78** |
| | | SD | | | | | 0.087 | 0.028 | 0.053 | 0.029 | 0.027 |
| | **0.2** | 11 | 344 | 437 | 123 | 182 | 0.65 | 0.78 | 0.44 | 0.93 | 0.72 |
| | | 12 | 294 | 431 | 97 | 189 | 0.61 | 0.82 | 0.44 | 0.93 | 0.72 |
| | | 13 | 304 | 451 | 110 | 223 | 0.58 | 0.80 | 0.39 | 0.94 | 0.69 |
| | | 14 | 270 | 522 | 207 | 93 | 0.74 | 0.72 | 0.44 | 0.93 | 0.73 |
| | | 15 | 306 | 483 | 147 | 170 | 0.64 | 0.77 | 0.41 | 0.95 | 0.71 |
| | | **mean** | | | | | **0.65** | **0.78** | **0.42** | **0.94** | **0.71** |
| | | SD | | | | | 0.063 | 0.039 | 0.020 | 0.010 | 0.012 |
